# Plastid phylogenomics of the *Sansevieria* clade (*Dracaena*; Asparagaceae) resolves a rapid evolutionary radiation

**DOI:** 10.1101/2021.01.05.421016

**Authors:** Iris van Kleinwee, Isabel Larridon, Toral Shah, Kenneth Bauters, Pieter Asselman, Paul Goetghebeur, Frederik Leliaert, Emily Veltjen

## Abstract

Sansevierias are a diverse group of flowering plants native to Africa, Madagascar, the Arabian Peninsula and the Indian subcontinent, popular outside their native range as low maintenance houseplants. Traditionally recognized as a distinct genus, *Sansevieria* was recently merged with the larger genus *Dracaena* based on molecular phylogenetic data. Within the *Sansevieria* clade, taxonomic uncertainties remain despite numerous attempts to classify the species. We aim to arrive at a robust phylogeny using a plastid phylogenomic approach, and estimate a time-frame of diversification to infer the evolutionary history of the group, including geographical and morphological evolution. Molecular data was obtained using genome skimming for 50 *Sansevieria*, representing all informal groups previously instated based on morphology, and two *Dracaena sensu stricto* species. The resulting Maximum Likelihood phylogenetic hypotheses are generally well supported, except for some very short branches along the backbone of the tree. The time-calibrated phylogeny indicates a recent rapid radiation with the main clades emerging in the Pliocene. Two well-supported clades align with previously defined informal groups, i.e., *Sansevieria* section *Dracomima*, characterised by the Dracomima-type inflorescence, and the *Zeylanica* group, native to the Indian subcontinent. Other morphologically defined informal groups are shown to be polyphyletic: a pattern due to convergent evolution of the identifying characters. Cylindrical leaves arose multiple times independently in the evolution of the *Sansevieria* clade and similarly, the Cephalantha-type inflorescence has originated multiple times from an ancestor with a Sansevieria-type inflorescence. To provide a more accessible tool for species identification and delimitation, genes and spacer regions were screened for variability and phylogenetic informativeness to investigate their potential as chloroplast DNA barcodes. Candidate chloroplast DNA barcodes include the *trnH-rpl12, ndhH-rps15, psbE-petL, psbT-psbN, rps18-rpl20* intergenic spacers, the chloroplast gene *rps8* and the first intron of *ycf3*.

## 1. Introduction

Sansevierias, a diverse group of flowering plants mostly found in dry habitats but also in wide variety of other habitats such as tropical forests and in coastal vegetation (Baldwin, 2016), are native to Africa, Madagascar, the Arabian Peninsula and the Indian subcontinent (Govaerts et al., 2020). Diverse sansevierias are found in many homes around the globe, popular because they are low maintenance houseplants. Common names linked to sansevierias are: ‘Mother-in-law’s tongues’, ‘Snake plants’ and ‘Bow string hemps’. A fair number of species are valued for their medicinal and ethnobotanical purposes (Khalumba et al., 2005; Haldar et al., 2010a, b; Takawira-Nyenya et al., 2014; Halyna et al., 2017; Maheshwari et al., 2017). Despite their economic importance, taxonomic uncertainty in terms of species identification and delimitation has resulted in a lack of progress in studying their evolution, diversity and ecology, and in assessing their conservation status.

Until recently, sansevierias were recognized as a distinct genus: *Sansevieria* Thunb. (e.g. Jankalski, 2008). However, molecular phylogenetic studies (Bogler and Simpson, 1996; Chen et al., 2013; Lu and Morden, 2014; Baldwin and Webb, 2016; Takawira-Nyenya et al., 2018) have shown that it is nested in the large genus *Dracaena* Vand. ex L., and consequently, it was placed in synonymy of the latter (Christenhusz et al., 2018; Takawira-Nyenya et al., 2018). *Dracaena* (190 species, Govaerts et al., 2020) is currently placed in Asparagaceae subfamily Nolinoideae (APG III, 2009; Kim et al., 2010; Chen et al., 2013; Lu and Morden, 2014; APG IV, 2016). Throughout this paper, the terms *Sansevieria* or sansevierias are used to describe the monophyletic group of *Dracaena* species (when excluding *Dracaena sambiranensis* (H.Perrier) Byng & Christenh.) that was formerly known as the genus *Sansevieria*, the term *Dracaena* to describe all other dracaenas, formerly placed in the genus *Dracaena*, and the term *Dracaena sensu lato* to refer to the newly circumscribed genus, including all the species belonging to the former genera *Chrysodracon* P.L.Lu & Morden, *Dracaena, Pleomele* Salisb., and *Sansevieria*.

The species of *Dracaena sensu lato* are united by similarities in floral characters and by 1–3 seeded berries (Mwachala and Mbugua, 2007). Within *Dracaena sensu lato*, sansevierias can be distinguished by a combination of morphological features, including fleshy, genuine succulent leaves, a herbaceous habit with rhizomes, and (mostly) unbranched thyrsose racemes (Table 1). Other members of *Dracaena sensu lato* generally lack (genuine) succulent leaves, can be trees and have (mostly) branched paniculate inflorescences (Table 1).

**Table 1.** Distribution and morphology of the (former) genera of the *Dracaena sensu lato*. Information is summarized from Mwachala & Mbugua (2007): *Sansevieria* and *Dracaena*, Brown (1914): *Pleomele*, Jankalski (2015) and Lu & Morden (2014): *Chrysodracon*.

|  | *Sansevieria* | *Dracaena* | *Pleomele* | *Chrysodracon* |
| --- | --- | --- | --- | --- |
| **Distribution** | Africa, Madagascar and South-East Asia. | Africa, southern Asia through to northern Australia, two species in tropical Central America and Cuba (Zona et al., 2014). | Central tropical Africa, Southern Asia and Malaysia. | Hawaii. |
| **Habit** | Herbaceous. | Herbaceous or tree forming. |  | Tree forming. |
| **Rhizome** | Strongly rhizomatous; rhizomes can be subterranean, on surface or aerial. | No rhizome |  |  |
| **Leaf succulence** | Genuine succulent and fleshy = xeromorphic leaves. | From thin leaves to sub-fleshy. No genuine succulent = mesomorphic leaves. |  | Never fleshy. |
| **Inflorescence** | (Mostly) unbranched thyrsose racemes. | (Mostly) branched paniculate |  | Paniculate. |
| **Flowering** | Nocturnally fragrant. |  |  | Diurnally fragrant. |
| **Perianth tube** | Perianth tube length variable. | Very short perianth tube with tepals divided to the base of the flower (Wagner et al., 1990). | Tepals connate for at least one-third of the perianth length (Wagner et al., 1990). | Tepals connate into a well-developed tube half to three-thirds the length of the perianth. |
| **Pollen** | Monosulcate and monoulcerate (Klimko et al., 2017). | Monosulcate and monoulcerate (Klimko et al., 2018). |  | Monoulcerate (Klimko et al., 2018). |

Within *Sansevieria*, three groups have been traditionally recognised based on inflorescence type (Newton, 2001; Mwachala and Mbugua, 2007; Jankalski, 2008, 2009; Mansfeld, 2015). In 2009, Jankalski recognised the three groups at sectional level, and later further subdivided them into 16 informal groups based on morphology (Jankalski, 2015). Molecular studies published to this date have not been able to draw strong conclusions about the evolutionary relationships between *Sansevieria* species (Bogler and Simpson; 1996; Chen et al., 2013; Lu and Morden, 2014; Baldwin and Webb, 2016; Takawira-Nyenya et al., 2018; Table 2). This because limited sampling of DNA regions and species resulted in low resolution. In addition, there have been questions about the reliability of species identification of the accessions sequenced. However, the most recent study by Takawira-Nyenya et al. (2018) showed that many of the morphology-based *Sansevieria* sections and informal groups appear to be para- or polyphyletic.

**Table 2.** Overview of Sanger sequencing studies which include *Sansevieria* and *Dracaena* species (number of species indicated with #).

| Reference | # *Dracaena* | # *Sansevieria* | Molecular markers |
| --- | --- | --- | --- |
| Takawira-Nyenya et al. (2018) | 19 | 26 | Chloroplast: *trnL* intron, *trnL-trnF* intergenic spacer and *rps16* intron; and the low-copy nuclear region At103 |
| Baldwin and Webb (2016) | 2 | 73 | Chloroplast intergenic spacers: between *trnT*, *trnL*, *trnF* |
| Lu and Morden (2014) | 31 | 34 | Chloroplast intergenic spacers: *trnL-trnF*, *ndhF-rpl32*, *trnQ-rps16*, and *rpl32-trnL* |
| Chen et al. (2013) | 4 | 1 | 4 chloroplast genes: *matK*, *rbcL*, *atpB*, *ndhF* |
| Bogler and Simpson (1996) | 1 | 1 | 2 nuclear genes: ITS1 and ITS2 |

Currently, *Sansevieria* comprises c. 80 species (Govaerts et al., 2020), listed in Appendix A. However, species delimitation has been a matter of discussion. As in most plant groups, morphology-based species delimitation in *Sansevieria* largely relies on floral characters (Brown, 1915; Jankalski, 2007; Mwachala and Mbugua, 2007). Few vegetative characters, such as leaf shape, leaf texture and margin colour, provide data to differentiate between species (Jankalski, 2015). This is because the leaves on a single *Sansevieria* plant may vary considerably and individual leaves also vary in morphology depending on the amount of shrivelling due to drought or age (Brown, 1915). To find alternative diagnostic characters, several studies investigated the informativeness of micromorphological characters to distinguish *Sansevieria* species, such as stomatal depth and cuticle thickness (Koller and Rost, 1988), pollen morphology (Klimko et al., 2017), and cell wall bands (Koller and Rost, 1988; van Kleinwee, 2018), which had varying, but generally unsatisfactory success.

One of the main problems hindering taxonomic revision of *Sansevieria* is that type material is not up to the required standards due to multiple reasons. The first reason is that only about 75% of species currently have a type specimen in a herbarium collection (Appendix A). The second reason is badly preserved and/or collected type specimens. A third reason is incomplete documentation: 13 described species have no type locality detailed below country level and for seven *Sansevieria* species even the country of origin is unknown (Appendix A). A fourth reason is type specimens described from cultivated plants with unknown wild origin: *D. longiflora* (Sims) Byng & Christenh., *D. trifasciata* (Prain) Mabb. and *D. zebra* Byng & Christenh. (former *Sansevieria metallica* Gérôme & Labroy), which invokes the possibility of the species being cultivars, hybrids, divergent growth forms of previously described species due to *ex situ* conditions,…. Other than lack of (good) typifications hampering taxonomy and identification; some species delimitations are doubtful given the incomplete or indistinct descriptions of the species.

The complex taxonomy of *Sansevieria* species results in very fragmented knowledge of population size and conservation status. Just a single *Sansevieria* species has been assessed using IUCN Red List criteria (Osborne et al., 2019; IUCN, 2020), despite the fact that many species are suffering from habitat destruction and/or are only known from a single location, and therefore are likely to be threatened like many other succulent plant groups (e.g., Larridon et al., 2014; Goettsch et al., 2015). For example, habitat destruction and overexploitation by local communities, mainly for fibre and medicinal use, causes *Sansevieria* species to be threatened in Zimbabwe (Takawira and Nordal, 2002). In Kenya, the centre of diversity of the group, home to 25 *Sansevieria* species, Newton (2018) noted that several species have gone extinct locally, including three species from their type localities, although they still occur elsewhere. According to The Red List of South African Plants (Raimondo et al., 2009), *D. zebra* (former *S. metallica*) is critically rare, only known from a single location. Other species only known from a single location are e.g., *D. pinguicula* (P.R.O.Bally) Byng & Christenh., *D. nitida* (Chahin.) Byng & Christenh., *D. longistyla* (la Croix) Byng & Christenh., *D. bugandana* Byng & Christenh., or even a single type collection e.g., *D. pedicellata* (la Croix) Byng & Christenh.

The aim of this study is to reconstruct the evolutionary relationships among Sansevierias using plastid phylogenomics and infer timing of diversification. This is the first well-sampled genome-scale phylogeny of *Sansevieria*, which provides new insights into the evolutionary history, including geographical and morphological evolution, and taxonomy of the group. Using the obtained genome-scale data, regions with potential chloroplast DNA barcodes (Hollingsworth, 2011) are identified. This study furthers our understanding of *Sansevieria*, which may benefit taxonomical and applied research, and conservation efforts.

## 2. Materials and methods

### 2.1 Plant material and DNA extraction

Leaf samples of *Sansevieria* species were collected from various botanic gardens and private collections. In total 52 samples were included of which 50 sansevierias, representing 46 species, and two *Dracaena* species (Appendix B). Collected leaf samples were selected to be as reliably identified as possible (i.e. collections which have been (largely) verified by experts) in combination with having at least two representatives of each of the informal groups defined by Jankalski (2015). Photographs of the accessions were acquired to verify whether their morphology falls within that of the informal group of Jankalski (2015) linked to the identification of the accession (Appendix B), but full verification was not always possible without inflorescence or other visible diagnostic characters. The identifications of accessions in the *Suffruticosa* group were verified using the key of Jankalski (2007). For other accessions, their morphology was compared with species descriptions. Multiple representatives of four species were included in the analysis (i.e. of *D. dooneri, D. parva, D. serpenta* Byng & Christenh. and *D. suffruticosa* (N.E.Br.) Byng & Christenh.) to evaluate intraspecific genetic variation or the suspicion of cryptic species. One *Nolina* Michx. species from GenBank (GenBank accession number: KX931462; McKain et al., 2016) was added to the dataset as outgroup.

Leaf samples were dried in silica-gel. The drying process of the fibrous, succulent leaf material was optimal when using thin, smaller pieces (± 0.5 × 1.5 cm) of the outermost green photosynthetic tissue, yielding high quality DNA extractions. The dried leaf samples were pulverized using BeadBeater (BioSpec, Oklahoma, USA). DNA was extracted following the protocol of Larridon et al. (2015) which is a modified CetylTrimethylAmmoniumBromide (CTAB) protocol (Doyle and Doyle, 1990) combined with a MagAttract suspension G (QIAGEN, Hilden, Germany) purification step (Xin and Chen, 2012). The protocol of Larridon et al. (2015) was altered by using 70% ethanol for cleaning instead of washing buffer in the final purification steps, which yielded higher DNA quality values. DNA quality control was executed using a NanoDrop 1000 Spectrophotometer (Thermo Fisher Scientific Inc., Waltham, MA, USA). Only samples with both optical density (OD) ratios of 280/260 and 260/230 higher than 1.80 were included in the Next Generation Sequencing (NGS) library. DNA samples were quantified with a Qubit 2.0 Fluorometer for which the Qubit dsDNA broad range (BR) Assay Kit was used (Life Technologies, Carlsbad, California, USA).

### 2.2 Library preparation, sequencing and data integrity

DNA samples were normalized to 0.75 ng. Library preparation was executed using the Illumina Nextera XT DNA Library Prep kit (Illumina Inc., California, USA). The sample libraries were validated by running 1 µL of undiluted library on an Agilent 2100 Bioanalyzer (Agilent Technologies, Palo Alto, California, USA) using a High Sensitivity DNA chip. The 52 sample libraries were subjected to standard normalization for which quantification was executed with the Qubit dsDNA high sensitivity (HS) Assay Kit and Qubit 2.0 Fluorometer (Life Technologies). Dilutions were performed using an EB-Tween solution (EB: Elution buffer) containing 10 mM Tris with 0.01% Tween at pH 8.0. The 52 normalized sample libraries were manually pooled into one pooled library, whereby each sample had a concentration of 7.5 nM, on a total volume of 200 µL. Pooling volumes were calculated using the Pooling Calculator (Illumina; https://support.illumina.com/help/pooling-calculator/pooling-calculator.html).

High-throughput sequencing using Illumina HiSeq 4000 (Illumina Inc.), was outsourced to Edinburgh Genomics (The University of Edinburgh, Edinburgh, Scotland). Reads were de-multiplexed by Edinburgh Genomics. Quality of the reads was inspected with FastQC version 0.11.3 (Andrews et al., 2011). Nextera adapter sequences (CTGTCTCTTATACACATCT) were trimmed using Cutadapt version 1.3 (Martin, 2011).

### 2.3 De novo assembly and mapping to reference

One *de novo* assembly of full chloroplast genome was executed for sample SA37B: *Dracaena conspicua* (N.E.Br.) Byng & Christenh. (Appendix B), which contained the highest number of reads (Appendix C). Contigs were generated by *de novo* assembly in QIAGEN CLC Genomics Workbench v.10.0.1 and Velvet v.1.2.10. In QIAGEN CLC Genomics Workbench the contigs were generated with an automatic word and bubble size and a minimum contig length of 200 base pairs. They were then exported with a threshold value of 20 and imported in Geneious v.8.1.9. (Kearse et al., 2012). Here, the contigs containing chloroplast genes were extracted the using Basic local alignment search tool (BLAST) (Altschul et al., 1990) to search for the CDSs (Coding DNA Sequences) of *Nolina atopocarpa* Bartlett against the different *de novo* assembled contigs. *Nolina atopocarpa* (Asparagaceae subfamily Nolinoideae) was used because it is the closest relative of *Sansevieria* (Kim et al., 2010), of which a fully annotated chloroplast genome was available (GenBank accession number: KX931462; McKain et al., 2016).

To reconstruct the chloroplast genome *de novo*, scaffolds were constructed based on the chloroplast contigs from QIAGEN CLC Genomics Workbench assembly and Velvet assembly (kmer size set to 91 and default settings) using the Geneious *de novo* assembler with medium sensitivity and default parameters. This resulted in two large scaffolds. The reads were again mapped to these scaffolds in Geneious (medium sensitivity, iterate 3 times), which resulted in an overlap between the two scaffolds. Because of sufficient overlap in the reads, the inverted repeat regions could also be fitted in the whole chloroplast genome sequence. All reads were once again mapped for additional verification. The constructed chloroplast was annotated from the *Nolina atopocarpa* chloroplast genome using the Live Annotate & Predict function in Geneious with similarity set to 75%. The annotated genome was aligned with the *Nolina atopocarpa* chloroplast genome using the Mauve algorithm with default settings in Geneious. The alignment was visually inspected for any bad or missing annotation transfers. The constructed chloroplast with annotations was visualized with OGDRAW (Lohse et al., 2007).

For the other samples, we attempted *de novo* assembly as described above, however due to ambiguous regions with low coverage, it was not possible to obtain the full chloroplast genome with high confidence. Hence, the chloroplast sequences of the other 51 samples were obtained by performing Map to Reference (default settings) to the chloroplast of SA37B in QIAGEN CLC Genomics Workbench and exported with a low coverage definition threshold value of 20, inserting N-ambiguities by low coverage, vote by conflict resolution and use of quality score. We acknowledge that possible rearrangements in the different chloroplast genomes would remain unnoticed, yet with the aim to construct a phylogenetic hypothesis, the result are not compromised.

### 2.4 Phylogenetic analyses and divergence date estimates

The 53 chloroplast genome sequences were aligned using the MAFFT tool using the CIPRES Science Gateway (Miller et al., 2015) and annotated using the *Nolina* reference genome (length of alignment: 162 166 bp). The alignment was trimmed using the heuristic automated1 algorithm in Trimal (Capella-Gutiérrez et al., 2009), resulting in a final alignment with a length of 159 637 bp provided in Appendix D. *Nolina* annotations were used to define blocks with coding, non-coding, inverted repeat and single copy regions.

The chloroplast genome alignment was analysed using a Maximum Likelihood (ML) approach in two ways, based on (1) an unpartitioned dataset, and (2) a dataset partitioned in 264 partitions. In both analyses one of the two inverted repeats was excluded because a) the two inverted repeat regions in the *de novo* assembly of SA37B were identical and b) the chloroplast genomes of the other 51 samples were constructed with map to reference, which does not allow discrimination between the two inverted repeats. The deleted region representing a copy of the Inverted Repeat lies between 46558–72292 in the alignment given in Appendix D, which is all the DNA between the CDS of *ycf1* and *psbA*. The 264 partitions represent 93 CDS partitions, the annotated gene: *infA*, 130 intergenetic spacers (IGS), 4 rRNA and 36 tRNA partitions. The 93 CDS partitions represent the 78 CDSs of *Nolina* (Appendix C) with their intervals (i.e. *atpF*: 2 exons, *clpP*: 3 exons, *ndhA*: 2 exons, *ndhB*; 2 exons, *petB*: 2 exons, *petD*: 2 exons, *rpl16*: 2 exons, *rpl2*: 2 exons, *rpoC1*: 2 exons, *rps12*: 3 exons, *rps16*: 2 exons, *ycf3*: 3 exons), excluding the seven CDSs present on the second copy of the inverted repeat. The CDS *rps12* has two exons on the inverted repeats (exon 2 and exon 3) and one exon outside the inverted repeat (exon 1). In the KX931462 *Nolina atopocarpa* chloroplast genome, some of the annotated CDS regions overlapped: *psbD* and *psbC* (overlap of 53 bp), *ndhK* and *ndhC* (overlap of 121 bp); and *atpE* and *atpB* (overlap of 4 bp). For the partitioning *psbD, ndhK* and *atpE* CDSs were not appointed in full, while *psbC, ndhC* and *atpB* were kept in their full length, making the genes adjacent instead of overlapping. The ML analyses were executed in IQ-TREE v1.7-beta18 (Nguyen et al., 2015). The unpartitioned analysis was set to run with 1000 ultrafast bootstraps using the “-bb” option, with no specification of a specific model resulting in IQ-TREE selecting the most optimal model for the data. The partitioned analysis was performed using the GTR model and with the “-spp partition_file”, “-bb” and “-bsam GENESITE” options in which each partition has its own evolutionary rate (Gadagkar et al., 2005; Chernomor et al., 2016). The partitioned analysis was set to run with 1000 ultrafast bootstraps, where IQ-TREE will resample the sites within partitions (i.e., the bootstrap replicates are generated per partition separately and then concatenated together).

For the dating analysis the inferred tree topology from the partitioned analysis was used, and branch lengths and node values were removed. Node calibration was implemented in R (v3.6.3), using the “estimateBound” function in the MCMCtreeR package (Puttnick, 2019). This estimates a uniform distribution across the hard minimum and soft maximum time constraints with 2.5% tail distributions (Puttick, 2019). The following secondary calibration points were used from Chen et al. (2013): (1) node *Dracaena* / *Sansevieria* (root *Sansevieria*): 3 mya (1–5 mya), and (2) node *Nolina* with *Dracaena* + *Sansevieria* (root *Dracaena*): 7.9 mya (2.5–11 mya). The calibrated tree was then dated using MCMCTree (dos Reis & Yang, 2019) in the PAML v4.9e package (Yang, 2007).

### 2.5 Classification, distribution and morphology

Classification data for each species identified for the accession was assembled, summarizing the main group (*Sansevieria* or *Dracaena*) and the 16 informal groups as published in Jankalski (2015). Distribution data for each species identified for the accession was assembled from literature, tabulating their distribution range and native TDWG (Taxonomic Databases Working Group) areas (Govaerts et al., 2020). To evaluate the morphology-based classification, two key characters used in the morphological classification were mapped on the resulting phylogenetic tree, namely: inflorescence type (Sansevieria-type, Dracomima-type and Cephalantha-type) and cylindrical *versus* flat leaves of adult plants. Appendix B provides the classification, distribution and morphological data from literature for the 52 accessions.

### 2.6 Screening for potential barcodes

The number of variable and parsimony informative sites were calculated for each contig alignment using AMAS (Borowiec, 2016). Partitions between 200–1000 bp in length, for which less than 5% of data was missing, were considered as potential chloroplast DNA barcodes. Of this selection of partitions, the top 5 variable sites and the top 5 parsimonious informative sites were highlighted. The list of markers was manually verified if they were reliably aligned.

## 3. Results

### 3.1 Sampling

*Dracaena powellii* (N.E.Br.) Byng & Christenh. was included in Jankalski’s classification (Jankalski, 2015) as a hybrid under the section *Dracomima*. He therefore did not indicate an informal group for this species. Based on the spirally-twisted leaves on an erect stem, the informal group of the species was noted in Appendix B as *Arborescens*. Other than a compilation of the metadata from literature, Appendix B includes newly generated data in the form of verification of informal groups and species identifications of the used accessions based on morphological data: 36/50 *Sansevieria* accessions were classified to have no indication for misidentification; 10/50 *Sansevieria* accessions had a suspicion of misidentification at the species level but not at the level of informal group; and 4/50 *Sansevieria* accessions were highlighted because there was a suspicion of misidentification even at the level of informal group. The morphology of the *Dracaena* species was not revised because they serve as outgroup only.

### 3.2 Sequence data

The high throughput sequencing (HTS) of the pooled library rendered 1,345,365 to 18,979,153 reads per accession, with an average of 6,754,755 reads (Appendix C).

### 3.3 Chloroplast genomes

The chloroplast genome (cp genome) of SA37B depicted in Figure 1, is 154,768 base pairs (bp) long. Almost all annotations from the *Nolina* chloroplast genome were transferred using the 75% similarity threshold, whereby the missing genes and CDSs (i.e. *petB, petD* and *rpl16*) where annotated during visual inspection of the genome alignment. The gene order of the SA37B chloroplast genome and the *Nolina* chloroplast genome is identical. The SA37B chloroplast genome can be found on GenBank with the accession number MW353256.

**Fig 1.**
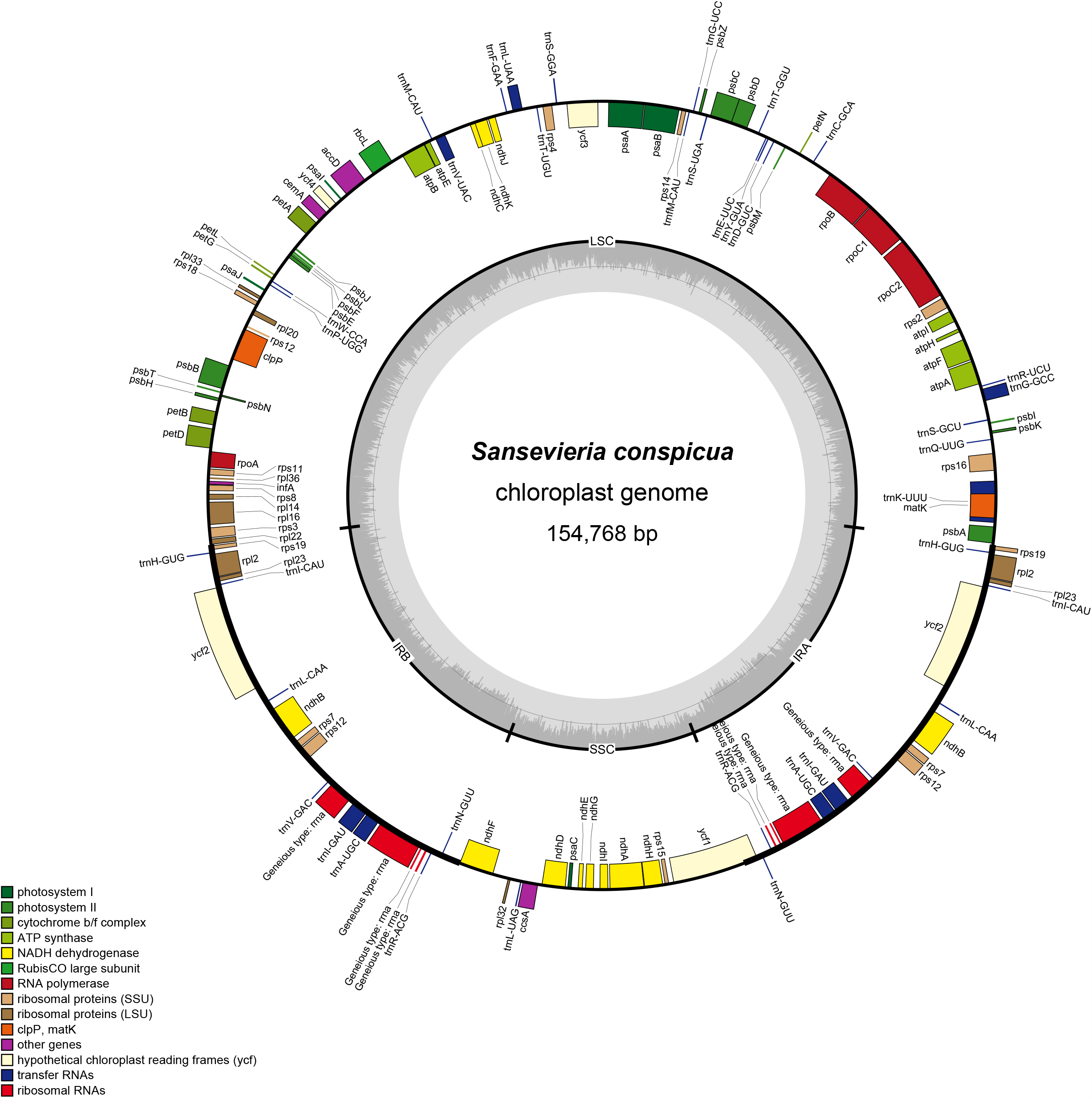
Full chloroplast genome of SA37B (*Dracaena conspicua* (N.E.Br.) Byng & Christenh.) annotated using *Nolina atopocarpa* Bartlett (NC_032708). Inverted Repeat (IR) regions are indicated, as well as the Short Single Copy (SSC) and Long Single Copy (LSC) regions. The circle inside the GC content graph marks the 50% threshold. GenBank accession number: MW353256.

Appendix C summarizes how many of the 85 chloroplast CDSs as identified in the chloroplast genome of *Nolina atopocarpa* (NC_032708) were retrieved in the 52 sequenced *Dracaena* and *Sansevieria* samples. The lowest number (57) of CDSs was found in the read data of *Dracaena stuckyi* (God.-Leb.) Byng & Christenh. For 28 of the 52 samples, all 85 CDSs identified in *Nolina atopocarpa*, were retrieved.

### 3.4 Phylogenetic hypothesis, divergence times, distribution and morphology

The ML phylogenetic tree based on the unpartitioned dataset is depicted in Appendix E. A dated ML phylogenetic tree based on the partitioned dataset is depicted in Figure 2, on which the evaluated geographic ranges and morphological characters from Appendix B are visualised. The same dated ML phylogenetic tree as in Figure 2 is given in Appendix F with the addition of the 95% confidence intervals on the nodes, which are tabulated explicitly next to the phylogenetic tree.

**Fig 2.**
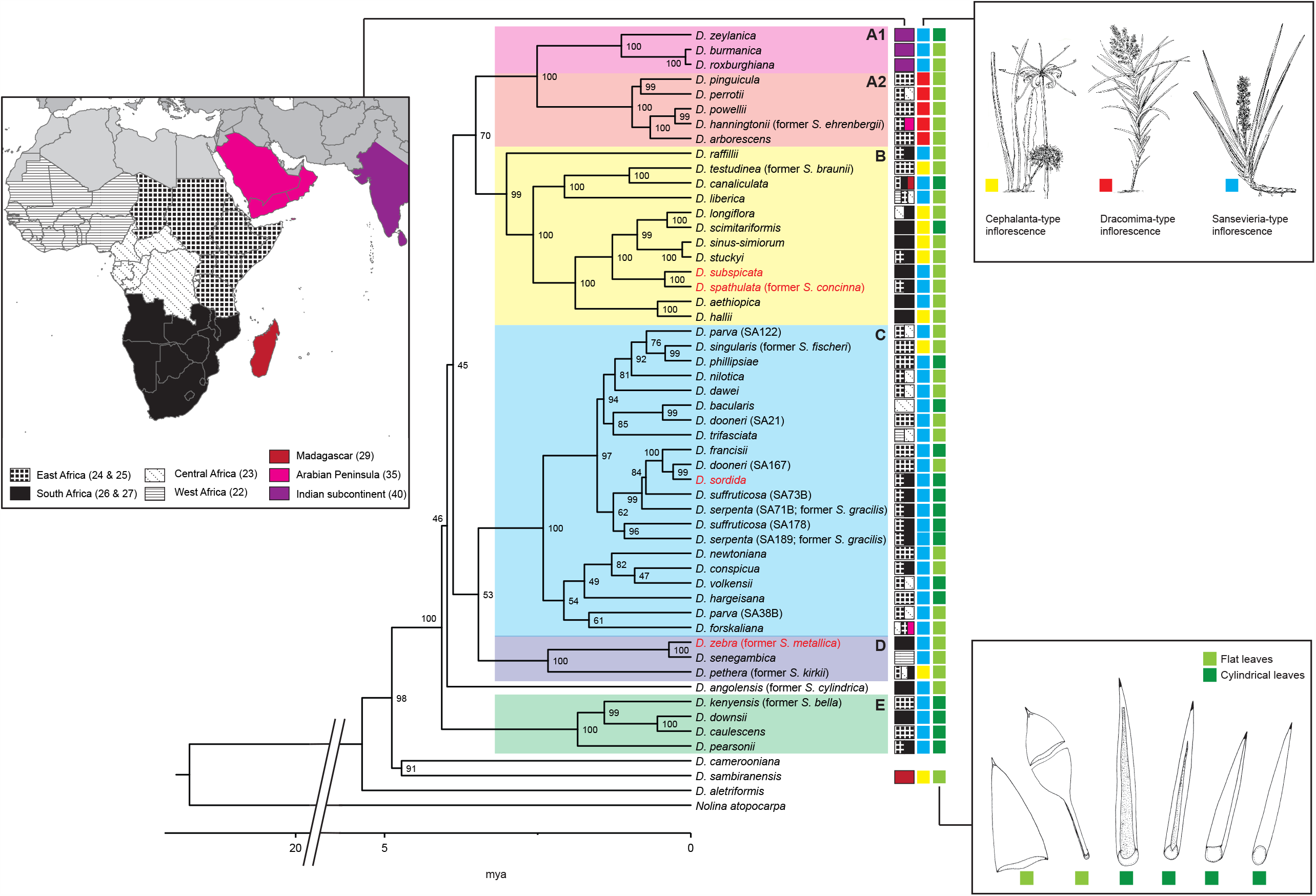
Dated phylogenetic tree inferred from the partitioned dataset of 50 *Sansevieria* species, two *Dracaena* species and *Nolina atopocarpa*. Branch lengths depict time, expressed in million years ago (mya). Bootstrap values are given above the branches. Species names in red indicate doubtful identifications (Appendix B). Geographic distinction is made according to the native Taxonomic Databases Working Group (TDWG) areas recorded for the species (Govaerts et al., 2020). Three inflorescence types were evaluated, illustrated by three examples from the monograph of Brown (1915): 1) the Cephalantha-type inflorescence represented by *Dracaena pethera* Byng & Christenh. (the former *S. kirkii* Baker), which consists of a congested unbranched pseudocapitate thyrsose raceme to umbelliform subcapitate on an elongate to subsessile scape; 2) the Dracomima-type inflorescence represented by *Dracaena powellii* (N.E.Br.) Byng & Christenh., which consists of elongate paniculate branched thyrsose racemes; and 3) the Sansevieria-type inflorescence, represented by *Dracaena suffruticosa* (N.E.Br.) Byng & Christenh., which consists of an elongate unbranched thyrsose raceme with flowers in interrupted cymose fascicles. Two adult leaf types were evaluated: flat *versus* cylindrical following the discrimination made in the given figure, adapted from Koller and Rost, 1988).

There are five main well-supported clades: Clade A–E (Figure 2, Appendix E, Appendix F). The relationships between the five clades have low support values and short branches. The relationships between the accessions within the clades are well-supported in clade A, B, D and E, while in clade C bootstrap values range from low to high support. The topologies of the unpartitioned (Appendix E) and the partitioned (Figure 2) phylogenetic tree are identical, with the exception of some of the relationships within clade C. Similarly, the bootstrap values for the well-supported clades are all 99 or 100 in both analyses, and more variable in de C clade. For the four species that had more than one accession in the analysis: *D. dooneri, D. parva, D. serpenta* and *D. suffructcosa*; no supported sister relationships were retrieved.

After the split from *Dracaena* the branches of the phylogenetic hypothesis are very short with time estimates between 5.188 and 2.003 mya (Appendix F: nodes 4–7 & 26), defining the stem nodes of five main *Sansevieria* clades (i.e. clade A–E). The five well-supported *Sansevieria* clades have their crown nodes estimated between 4.004 and 0.794 mya (Appendix F: nodes 8, 15, 27, 47 & 49).

### 3.5 DNA barcodes

Appendix G depicts summary statistics of the different genomic regions such as number and proportion of variable sites, number and proportion of parsimony informative sites and length. Regions between c. 200–1000 bp in length, for which less than 5% of data was missing in our dataset, that have a high proportion of parsimonious informative sites are indicated in yellow. The following regions are most promising as potential chloroplast DNA barcodes to identify *Sansevieria* species: the *trnH-rpl12, ndhH-rps15, psbE-petL, psbT-psbN, rps18-rpl20* intergenic spacers, the chloroplast gene *rps8* and the first intron of *ycf3*.

## 4. Discussion

### 4.1. Sampling

Our study used 50 *Sansevieria* accessions that represent 46 of the c. 80 described species (Appendix B), whereby all the informal groups (Jankalski, 2015) are represented by two or more *ex situ* accessions that are reliably identified and verified by experts, and by the authors at least at the level of informal group (Appendix B). The relationships among these 46 *Sansevieria* species are studied, for the first time, on a genomic level which has rendered a high quantity of data and well-supported clades (Figure 2). The combination of this comprehensive sampling and substantially larger molecular dataset (Table 2) has resulted in a significant improvement of the knowledge on relationships among *Sansevieria* biodiversity. Although identification of some *Sansevieria* accessions is uncertain (Appendix B), our study enables a first robust phylogeny of a significant amount of morphological diversity currently present in important *ex situ* collections. It provides an evolutionary framework for the group to which can be expanded in future to study the evolution and diversity of the group in more detail. In particular, it would be valuable to add more wild-collected accessions, preferably from type locations. In all cases, morphology of the accessions should be compared to the original species descriptions; and in cases where the type material and species description are unclear or lacking: a revision and redocumentation of the species is advised like the work of Newton (2009) and Mansfeld & Gerhard (2015), executing typifications where necessary.

The lack of a sister relationship between the multiple accessions of single species included (i.e. *D. dooneri, D. parva, D. serpenta* and *D. suffructicosa*) can be due to misidentifications or the presence of cryptic species. Identification of the accessions should be revised, and the species are here highlighted for taxonomic revision. This illustrates the interactive nature between phylogenetic studies and taxonomy, rather than a phylogenetic hypothesis being a final product. The SA167 *D. dooneri* accession was collected in Kenya, which is the type country, while for the SA21 *D. dooneri* sample, the collecting locality is unknown (Appendix B) – yet the specimen morphology matched the species description more closely. The SA122 *D. parva* sample was collected in Uganda; and the SA38B sample was collected in Burundi (Appendix B), originally included to study the intraspecific variation of this species. The morphology of the two *D. parva* accessions does not appear to be divergent from expectations for the species. A possible explanation is that the accessions represent convergent evolution to the “grass-like” habit from different evolutionary trajectories.

### 4.2. Sequence data

The high variety in number of reads and the 24 samples in which not all CDSs identified in *Nolina* were retrieved (Appendix C) is most likely linked to the choice of the Nextera XT kit, which is designed for samples with low amounts of DNA and subsequently most of the samples had to be severely diluted, to fit the Nextera XT demand of maximally 1 ng DNA. The dilution invokes more room for human and/or pipetting errors, most likely leading to different start DNA quantities and hence finally different quantities in the pooled library per sample. As it was difficult to construct the full chloroplast genomes, we advise to pool a smaller number of samples in one sequencing lane. The transposase used in the Nextera system, as with any enzymatic system, could also have invoked a slight bias in the binding reaction, hampering complete retrieval of the chloroplast genome. Despite possible improvements to the lab methods used, it is unlikely that repeating the experiment will significantly improve phylogenetic signal. The aim of this study was not to retrieve 52 whole chloroplast genomes, but to find informative plastid genome regions able to differentiate *Sansevieria* species, which was successful as the majority of the chloroplast base pairs were recovered (Appendix C).

### 4.3 Chloroplast genomes

It is to be expected that chloroplast genomes from closely related land plants are conservative in their general structure (Palmer et al., 1988), although exceptions are known in angiosperms (Downie and Palmer, 1992; Wicke et al., 2011; Röschenbleck et al., 2017). Comparing the chloroplast genome of a *Sansevieria* species SA37B (*D. conspicua*) with *Nolina atopocarpa* confirms the conservative nature of the chloroplast genome. In further analyses, it is advised to run more *de novo* assemblies, to rule out gene rearrangements within the *Sansevieria* clade (e.g. Cauz-Santos, et al., 2020).

### 4.4 Phylogenetic hypothesis, divergence times, distribution and morphology

Up until now, Sanger sequencing-based methods were used to sequence only a small number of loci (Table 2), which represent a tiny fraction of the genomic information available in a plant cell. Earlier Sanger sequencing-based studies (Table 2) had issues with low resolution (e.g., Lu and Morden, 2014; Baldwin and Webb, 2016; Takawira-Nyenya et al., 2018). Although using the full chloroplast genome has improved the resolution and rendered new results including strong support for the main clades, unsupported deeper nodes (i.e. between clades A–E), as well as unsupported recent relationships remain (i.e. of a number of sister species within clade C). The lack of support for the deeper relationships are most likely a result of a rapid radiation with the main clades emerging in Pliocene (Figure 2, Appendix F), leaving little phylogenetic signal in the chloroplast genome to infer relationships among the main clades. The time interval of rapid *Sansevieria* evolution leading to the five main clades is estimated between 5.188 and 2.003 mya (Appendix F). This range overlaps with, but is overall younger than, the age ranges found of recent rapid radiations in other studied succulent groups, such as the Aizoaceae (8.7–3.8 mya; Klak et al., 2004), Cactaceae (10–5 mya; Arakaki et al., 2011) and *Agave sensu lato* (12.34–4.62 mya; Flores-Abreu et al., 2019). In the case of *Agave* (Flores-Abreu et al., 2019), the authors speculated that the rapid recent diversification could be attributed co-evolution with their pollinator community (Flores-Abreu et al., 2019), which for *Sansevieria* could also be a (co-)driving force for the rapid radiation. More research on the *Sansevieria* pollinator community could explore this possibility in more depth.

The lack of support for the most recent relationships are most likely caused by a) the high amount of vegetative propagation in comparison to sexual propagation resulting in a slow accumulation of phylogenetically informative mutations (Ma et al., 2017); b) recent speciation (Parks et al., 2009); c) important linking taxa that are missing from the analysis (Nabhan and Sarkar, 2012); and/or d) incorrect taxonomic splitting of species that still have gene flow.

As this study mainly focused on relationships within the *Sansevieria* clade, only two *Dracaena* species were included as outgroup. As a result, the phylogenetic tree (Figure 2) cannot serve as additional evidence for the *Dracaena* - *Sansevieria* relationship. However, it does confirm earlier studies that placed *D. sambiranensis* in *Dracaena* rather than in *Sansevieria* (Lu and Morden, 2014; Takawira-Nyenya et al., 2018). Our study also indicates a young age of the *Sansevieria* clade, which is estimated to have originated in the Late Miocene – Pliocene (c. 6.573–2.671 mya) (Appendix F: node 3).

Clade A (Figure 2) comprises two subclades of which one consist of sansevierias that colonised the Indian subcontinent (clade A1), classified by Jankalski as the *Zeylanica* group (Jankalski, 2015), and one clade consisting of sansevierias with the Dracomima-type inflorescence (clade A2), classified by Jankalski as section *Dracomima* (Jankalski, 2015). Assuming that the origin of *Sansevieria* lies in Africa, the monophyly of clade A1 supports a single colonisation event of *Sansevieria* to the Indian subcontinent between 3.565–0.431 mya (Appendix F: node 8 & 9). Since identification to species level in the *Zeylanica* group is difficult (Appendix B), addition of more accessions with wild origin data from the Indian subcontinent is recommended in future research. The two representatives of the *Ehrenbergii* group (Jankalski, 2015): *D. hanningtonii* Baker and *D. perrotii* (O.Warburg) Byng & Christenh. do not form a clade. This result indicates that the species delimitation within section *Dracomima* (Jankalski, 2015) based on spirally (*Arborescens* group) vs. distichously arranged leaves (*Ehrenbergii* group) has no evolutionary significance.

Clade B (Figure 2) is well-supported and includes species with either a Cephalantha-type or a Sansevieria-type inflorescence, most of which have flat leaves. As both inflorescence types do not form supported clades, the two linked sections (Jankalski, 2008, 2009) are not phylogenetically supported. Within clade B, all relationships are very well supported. We retrieved one clade that has species with a distribution centred in East Africa, and a second clade that comprises species with a distribution centred in southern Africa. The species of clade B have been classified in eight different informal groups (Appendix B) of which only one is supported in our results, i.e. the *Subspicata* group. However, when examining the accessions morphologically and geographically, doubt is casted on this support (Appendix B). Within clade B, one well-supported subclade comprises of all species with a Cephalantha-type inflorescence: *D. longiflora, D. scimitariformis* (D.J.Richards) Byng & Christenh., *D. sinus-simiorum* (Chahin.) Byng & Christenh. and *D. stuckyi* (God.-Leb.) Byng & Christenh. However, two of the four accessions need further verification of their identity (Appendix B).

Clade C (Figure 2) represents a well-supported main clade in *Sansevieria* composed of accessions placed in six informal groups (Appendix B), most with a Sansevieria-type inflorescence. It is composed of one well-supported subclade and 6 accessions with low support (Figure 2). Geography, more than morphology seems to be indicative for some of the found supported relationships within this clade. For example the distribution range of sister species *D. singularis* (N.E.Br.) Byng & Christenh. and *D. phillipsiae* (N.E.Br.) Byng & Christenh. overlap in Ethiopia and Somalia. Although both species have cylindrical leaves, their habit and floral morphology are very different. Similarly, *D. sordida* (N.E.Br.) Byng & Christenh., *D. francisii* (Chahin.) Byng & Christenh. and *D. dooneri* form a supported clade and all three accessions were collected from Kenya (Appendix B). The accession included of *D. forskaliana* was collected in Yemen (Appendix B) and forms an unsupported sister relationship with *D. parva* (SA38B) from Burundi dated 2.586–0.785 mya (Appendix F: node 46). Looking at the supported nodes only, this migration from Africa to the Arabian Peninsula is dated to be younger than 3.351 mya (Appendix F: node 27). Interestingly, the supported clade A2 representing species from section *Dracomima*, including *D. hanningtonii* with known distribution in the Arabian Peninsula, and the supported clade C, containing *D. forskaliana* with known distribution in the Arabian Peninsula, indicate two separate dispersal events of *Sansevieria* from Africa to the Arabian Peninsula.

Clade D (Figure 2) is well-supported and comprises species with flat leaves (Appendix B). It is not clear what unites these species at first, as they have been classified in two sections and three informal groups and their distribution encompasses West, Central, East and southern Africa. However, morphologically, the accession identified as *D. zebra* fits better in the *Trifasciata* informal group rather than the *Hyacinthoides* informal group (Appendix B), which reduces the clade to two sections and two informal groups. Geographically the two accessions of clade D for which the locality data in known, originate from two different countries: the Democratic Republic of the Congo and Senegal (Appendix B).

*Dracaena angolensis* (Welw. ex Carrière) Byng & Christenh., a species with a Sansevieria-type inflorescence and cylindrical leaves, does not fall within a supported clade.

Clade E (Figure 2) is another well-supported clade including species that have been classified in two informal groups: *Pearsonii* and *Suffruticosa*. The members of this clade all have a Sansevieria-type inflorescence and cylindrical leaves (Appendix B). The geography of the four samples included is known and is quite extensive: Namibia, Kenya, Zimbabwe and the Democratic Republic of Congo (Appendix B).

In general, the *Hyacinthoides* group of section *Sansevieria* is very heterogeneous in its macromorphology, and this is reflected in the phylogenetic positions of the species in the tree (Figure 2), as had already been suggested by the Sanger sequencing-based study of Takawira-Nyenya et al. (2018). The morphological characterization of this group was not expected to have any evolutionary value, because it mostly includes species that simply do not fit in any of the other groups (i.e. a ‘wastebasket taxon’).

### 4.5 Further notes on classification, ecology and morphology

The short phylogenetic distance between *Sansevieria* species and the easy occurrence of (artificial) hybridization (Pate et al., 1954; Menzel and Pate, 1960), invoke doubt about whether or not gene flow still occurs between some of the described species. For example, Pfennig (1979) suggested that *D. powellii* may be a natural hybrid of *D. perrotii* and *D. arborescens* (Cornu ex Gérôme & Labroy) Byng & Christenh., as it arose in regions where the two species are sympatric. There is no definite proof for his suggestion, as artificial hybrids of these two species are only 10 cm high and do not resemble *D. powellii*.

The reproductive ecology of sansevierias is poorly studied; we only know that they flower at night emitting a strong pleasant scent, attracting insects (possibly hawkmoths) for pollination (Tanowitz and Koehler, 1986). Studies on seed dispersal have not been conducted but are important to assess the geographic range in which species can reproduce. Although there is hardly any research on hybridization in the wild, fertile hybrids are easily formed in experimental settings (Pate et al., 1954; Menzel and Pate, 1960) and differ greatly in morphology. Consequently, it is likely that speciation by hybridization may occur regularly in *Sansevieria*. Studying reproduction ecology and hybridization in *Sansevieria* will also provide better insights into species boundaries.

Brown (1915) hypothesized that species with cylindrical leaves originated from ancestral forms having flattened leaves based on plant ontogeny (juvenile plant leaves are first flat to concave and later transition to cylindrical adult leaves). The cylindrical leaf-type represents a synapomorphy only for clade E, and further arose multiple times in *Sansevieria* (Figure 2). Cylindrical leaves may have evolved on multiple occasions supported by a relatively easy genetic ‘switch’, activated by similar selection pressures of the environment. In extreme drought and high solar irradiation conditions, cylindrical leaved Sansevierias have been suggested to be even more water efficient, than for example flat leaved *Dracaena* species (Sreenivasan et al., 2011).

### 4.6 Species identification through DNA barcoding

A DNA barcode needs to meet several criteria, namely they need to: a) contain a high proportion of phylogenetically informative sites; b) be short (400–800 base pairs), as to facilitate current capabilities of DNA extraction and amplification (Kress and Erickson, 2008); and c) be flanked by conservative regions so that universal Forward and Reverse primers can be designed (e.g. Ford et al., 2009; Hollingsworth et al., 2011).

None of the six listed barcodes have been used in previous Sanger sequencing studies of *Sansevieria* (Bogler and Simpson, 1996; Chen et al., 2013; Lu and Morden, 2014; Baldwin and Webb, 2016; Takawira-Nyenya et al., 2018). For the practical application of species identification, for example to identify plants in *ex situ* collections, the proposed barcodes can be sequenced in parallel with careful examination of their morphology. However, even used together they may not enable to differentiate between sister species, and genomic tools such as genome skimming (cf. this study) or target capture sequencing are more appropriate for species identification with the goal of further elucidating the taxonomy of *Sansevieria*. These techniques have become much cheaper (Hale et al., 2020), work well with herbarium material (Brewer et al., 2019), and universal enrichment panels have been shown to be able to resolve relationships in a range of plant groups (e.g., Fragoso-Martínez et al., 2017; Larridon et al., 2020; Shah et al., submitted), even between closely related species, reducing the need to develop more expensive taxon-specific custom enrichment panels.

## 5. Conclusions and further research

Our plastid phylogenomic analyses provide new insights into evolutionary relationships between *Sansevieria* species, and the link with geographical distribution and morphology. Although low support was retrieved for some nodes in the backbone of the *Sansevieria* clade, most clades and relationships between species are well-supported. *Dracaena sambiranensis* was positioned outside the clade comprised of former *Sansevieria* species. The time-calibrated phylogeny indicates a recent rapid radiation with the main clades emerging in the Pliocene. Within the *Sansevieria* clade, two of the well-supported clades clearly align with morphological groups previously defined by Jankalski (2015), i.e., *Sansevieria* section *Dracomima* and the *Zeylanica* group. Other sections and informal groups are shown to be polyphyletic. Cylindrical leaves have evolved multiple times in the evolution of the *Sansevieria* clade, hypothesised to be correlated to drought. Similarly, the Cephalantha-type inflorescence has originated multiple times from an ancestor with a Sansevieria-type inflorescence. For future studies, we recommend continuing to work with phylogenomic data given the low sequence divergence in the group, whereby universal targeted sequencing enrichment panels such as Angiosperms353 (Johnson et al., 2019) can be employed to further explore the potential of nuclear DNA in studying the evolutionary history *Sansevieria*. Ideally more accessions collected from type localities should be sequenced to support taxonomic revision of the group. Multiple accessions per species collected from throughout its distribution range should be included to study intra-versus interspecific genetic variation. Potential chloroplast DNA barcodes to quickly identify *Sansevieria* species at a lower cost are the *trnH-rpl12, ndhH-rps15, psbE-petL, psbT-psbN, rps18-rpl20* intergenic spacers, the chloroplast gene *rps8* and the first intron of *ycf3*.

## Acknowledgements

We thank the Friends of the Ghent University Botanical Garden for funding the molecular component of this study. For help with the NGS lab work, we thank Wim Baert of Meise Botanic Garden. For collection of samples, we acknowledge the collection managers of Meise Botanic Garden, Royal Botanic Gardens Kew, Ghent University Botanical Garden, University of Potsdam Botanical Garden, University of Heidelberg Botanical Garden, and Strasbourg University Botanical Garden. Hereby we would explicitly like to thank Paul Rees (Royal Botanic Gardens Kew), Michael Burkart (Potsdam Botanical Garden) and Anthony Beke (Strasbourg University Botanical Garden). We also received leaf samples from the private collections of Len Newton, Gilfrid Powys and Tom Forrest, who enthusiastically supported us with their knowledge of the *Sansevieria* diversity.

## Appendices

**Appendix A**. 79 described former *Sansevieria* species, now included in *Dracaena*, with their type localities and absence/presence data on their type collection. Accepted species names indicated in yellow are a new combination. Type locations in yellow are those considered to have no locality within the country. Type locations in red are those considered not to even have a type country linked to the species name.

**Appendix B**. Overview of the 52 accessions (50 *Sansevieria*, 2 *Dracaena*) used in final library for High-throughput sequencing, with the classification, distribution and morphology metadata linked to their species identification; as well as their appointed clade as retrieved from the dated, partitioned Maximum Likelihood phylogenetic hypothesis (see Figure 2). Main group: the former *Sansevieria* genus or the former *Dracaena* genus. Sections as recognised by Jankalski 2008, 2009: *Sansevieria, Dracomima* or *Cephalantha*; following the three inflorescence types: Cephalantha-type – consists of a congested unbranched pseudocapitate thyrsose raceme to umbelliform subcapitate on an elongate to subsessile scape; Dracomima-type – consists of elongate paniculate branched thyrsose racemes; and Sansevieria-type – consists of an elongate unbranched thyrsose raceme with flowers in interrupted cymose fascicles. Informal groups following Jankalski 2015. Geographical area: IND: Indian subcontinent; SA: South Africa; EA: East Africa: WA: West Africa; CA: Central Africa: MAD: Madagascar; ARAB: Arabian Peninsula (See Figure 2).

**Appendix C**. The 50 *Sansevieria* and 2 *Dracaena* samples obtained by high-throughput sequencing using Illumina HiSeq 4000 (Illumina Inc.) and an Illumina Nextera XT kit. Their number (#) of reads are indicated as well as the number (#) of chloroplast Coding DNA Sequences (CDS) found in contigs after stripping the alignment. The numbers in parenthesis indicate the genes duplicated in the inverted repeat regions. *Nolina atopocarpa* voucher: NC_032708, GenBank accession number: KX931462; McKain et al., 2016; was used as a reference genome.

**Appendix D**. Fasta-file containing the file alignment of the 52 sequenced accessions of *Sansevieria* and *Dracaena* (See Appendix B).

**Appendix E**. An unpartitioned Maximum Likelihood hypothesis of the 52 sequenced *Sansevieria* and *Dracaena* accessions. Numbers on the nodes represent bootstrap values. Clades correspond to the clades of Figure 2. *Nolina atopocarpa* Voucher: NC_032708, GenBank accession number: KX931462; McKain et al., 2016; was used as outgroup.

**Appendix F**. The same Maximum Likelihood tree of 52 accessions of *Sansevieria* and *Dracaena* (Appendix B) constructed from the partitioned analysed chloroplast genome alignment (Figure 2), whereby the 95% confidence intervals on the ages of the nodes is given in the table. Numbers on the nodes represent an identifier number for that specific node, to be matched with the 95% confidence intervals of the table. Node 2 and node 4 are the calibrated nodes.

**Appendix G**. The AMAS (Borowiec, 2016) statistics of the partitions of the chloroplast genome alignment (Appendix D) of the 52 *Sansevieria* and *Dracaena* accession from Appendix B. The chloroplast regions in yellow are considered to have the greatest potential to serve as chloroplast DNA barcodes.

