## Supplementary figures and images for "Plastid phylogenomics of the *Sansevieria* clade (*Dracaena*; Asparagaceae) resolves a rapid evolutionary radiation"

### Appendix E

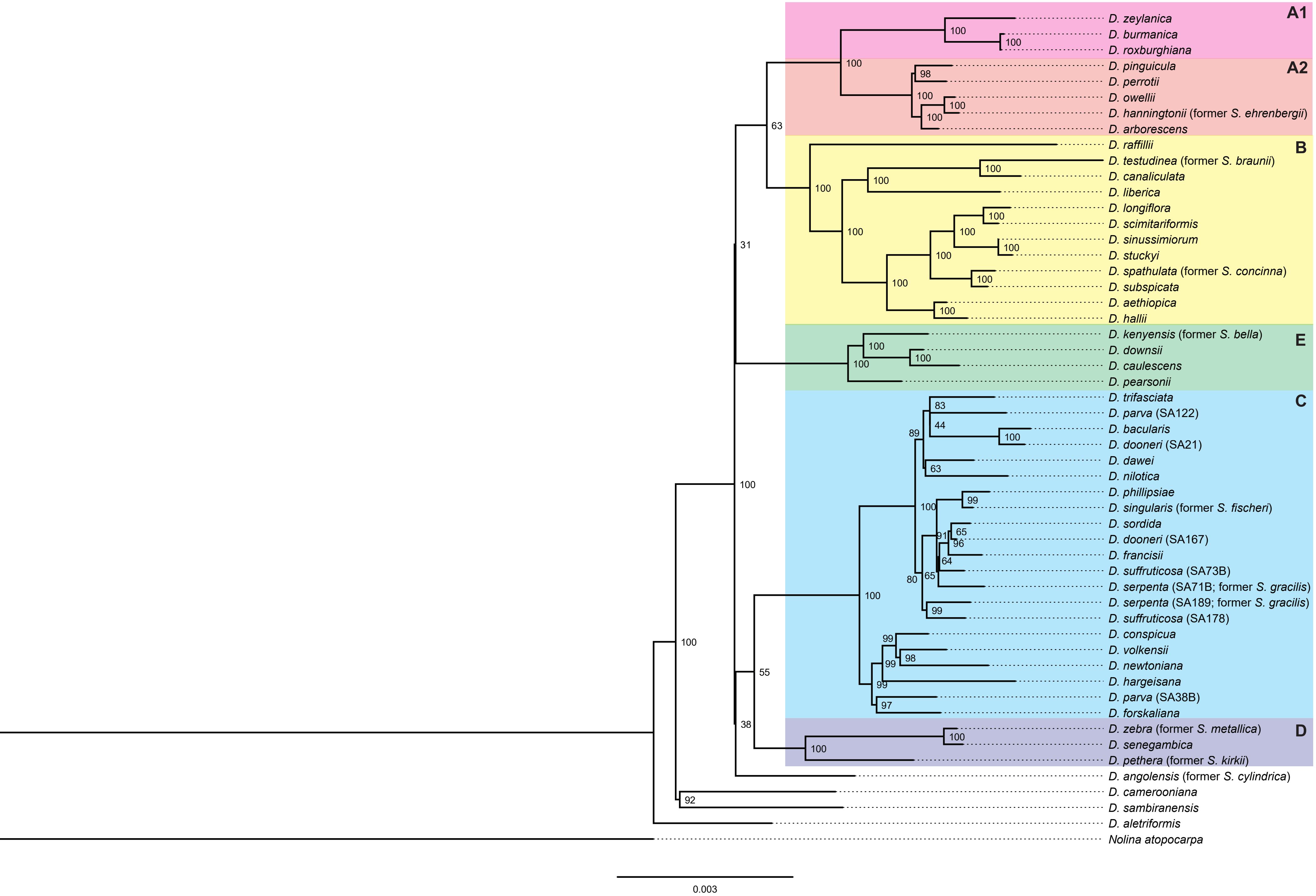

### Appendix F

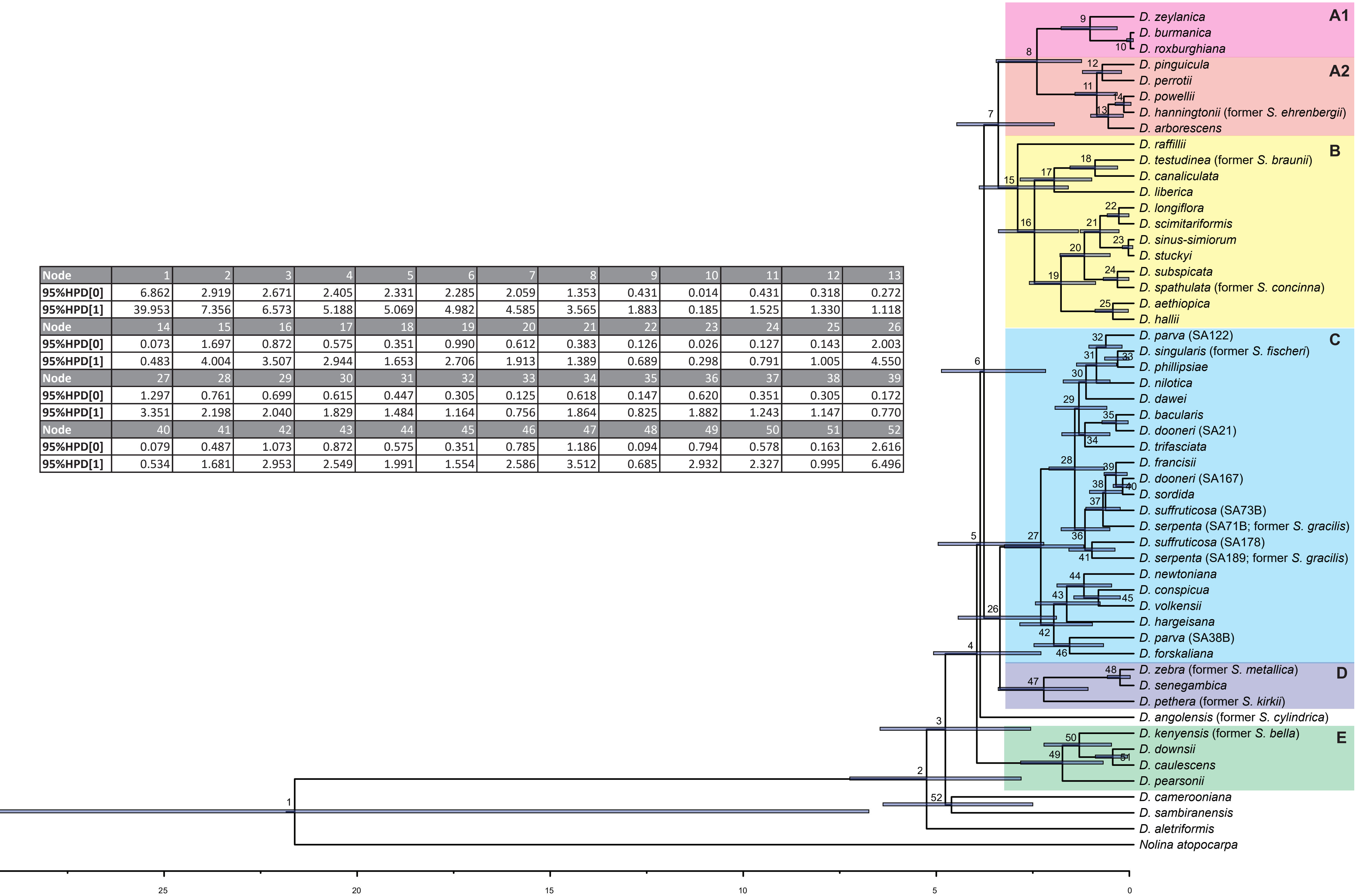


| Node | 1 | 2 | 3 | 4 | 5 | 6 | 7 | 8 | 9 | 10 | 11 | 12 | 13 |
|---|---|---|---|---|---|---|---|---|---|---|---|---|---|
| 95%HPD[0] | 6.862 | 2.919 | 2.671 | 2.405 | 2.331 | 2.285 | 2.059 | 1.353 | 0.431 | 0.014 | 0.431 | 0.318 | 0.272 |
| 95%HPD[1] | 39.953 | 7.356 | 6.573 | 5.188 | 5.069 | 4.982 | 4.585 | 3.565 | 1.883 | 0.185 | 1.525 | 1.330 | 1.118 |
| Node | 14 | 15 | 16 | 17 | 18 | 19 | 20 | 21 | 22 | 23 | 24 | 25 | 26 |
| 95%HPD[0] | 0.073 | 1.697 | 0.872 | 0.575 | 0.351 | 0.990 | 0.612 | 0.383 | 0.126 | 0.026 | 0.127 | 0.143 | 2.003 |
| 95%HPD[1] | 0.483 | 4.004 | 3.507 | 2.944 | 1.653 | 2.706 | 1.913 | 1.389 | 0.689 | 0.298 | 0.791 | 1.005 | 4.550 |
| Node | 27 | 28 | 29 | 30 | 31 | 32 | 33 | 34 | 35 | 36 | 37 | 38 | 39 |
| 95%HPD[0] | 1.297 | 0.761 | 0.699 | 0.615 | 0.447 | 0.305 | 0.125 | 0.618 | 0.147 | 0.620 | 0.351 | 0.305 | 0.172 |
| 95%HPD[1] | 3.351 | 2.198 | 2.040 | 1.829 | 1.484 | 1.164 | 0.756 | 1.864 | 0.825 | 1.882 | 1.243 | 1.147 | 0.770 |
| Node | 40 | 41 | 42 | 43 | 44 | 45 | 46 | 47 | 48 | 49 | 50 | 51 | 52 |
| 95%HPD[0] | 0.079 | 0.487 | 1.073 | 0.872 | 0.575 | 0.351 | 0.785 | 1.186 | 0.094 | 0.794 | 0.578 | 0.163 | 2.616 |
| 95%HPD[1] | 0.534 | 1.681 | 2.953 | 2.549 | 1.991 | 1.554 | 2.586 | 3.512 | 0.685 | 2.932 | 2.327 | 0.995 | 6.496 |
